# Glucocorticoid-sensitive period of corticotroph development – implications in early life stress

**DOI:** 10.1101/2022.06.13.495881

**Authors:** Guy Peles, Amrutha Swaminathan, Gil Levkowitz

**Author notes:** equal contribution (Guy Peles, Amrutha Swaminathan).

## Abstract

Corticotrophs are intermediaries in the hypothalamic-pituitary-adrenal (HPA) axis, which plays a crucial role in vertebrates’ stress response. The HPA axis displays an intricate mode of negative feedback regulation, whereby the peripheral effector, cortisol inhibits the secretion of its upstream regulator, adrenocorticotropic hormone (ACTH) from proopiomelanocortin (POMC)-expressing cells in the pituitary. While the feedback regulation of the HPA axis is well characterized in the adult organism, the effect of feedback regulation on the development of corticotrophs is poorly understood. Here, we study the effect of glucocorticoids on the development of POMC-expressing cells in the zebrafish pituitary. The development of POMC cells displayed striking robustness in terms of their steady increase in numbers between 2-6 days post fertilization. Inhibition of endogenous glucocorticoid synthesis resulted in an increase in POMC cell number due to reduced developmental feedback inhibition of cortisol on POMC cells. Conversely, addition of exogenous dexamethasone at a critical developmental window led to a decrease in POMC cell number, mimicking greater feedback control. Finally, developmental dysregulation of ACTH levels resulted in impaired anxiety-like and stress-coping behaviours. Hence, we identified a sensitive developmental window for the effect of glucocorticoids on corticotrophs and demonstrate the downstream effect on stress-responsive behaviour.

## Introduction

The hypothalamic-pituitary-adrenal (HPA) axis is a central neuroendocrine system, which allows all vertebrates to respond and adapt to physiological and psychological stressors in order to maintain homeostasis. Activation of the axis is achieved by the release of corticotropin-releasing hormone (CRH) from hypothalamic neurons thereby triggering the secretion of adrenocorticotropic hormone (ACTH) from pituitary cells to the general circulation to finally induce the secretion of corticosteroids from the adrenal cortex (1). Additionally, the HPA axis is regulated by a negative feedback loop to re-establish homeostasis, whereby corticosteroids have an inhibitory effect on the secretion (rapid response) and synthesis (delayed response) of CRH and ACTH in the hypothalamus and pituitary (2, 3). In addition to short-term alterations at the level of synthesis and secretion of these hormones, there is evidence of long-term changes in the volume and total cell mass (4, 5).

The HPA axis develops early during embryonic stages and is already fully functional at later gestational stages in mammals (6). This development is closely linked to early life experience, including parental influences like stress, maternal grooming and separation (6). Similarly, zebrafish have developed all components of the highly conserved HPA axis by 3-4 days post fertilization (7), and 5-6 day-old zebrafish embryos show robust stress responsive behaviour including rapid increase in cortisol levels following a stress challenge (8-10).

While the development of the HPA axis is understood, the dynamics of feedback regulation has not been explored in developmental stages, due to the challenges associated with inaccessibility of young mammalian embryos. Furthermore, while the effect of cortisol on the levels of circulating CRH and ACTH is well-documented, its effect on the physiology of the CRH-positive and ACTH-positive cells in the brain is less understood.

Here, we focus on the effect of changes in glucocorticoid levels on the development of corticotrophs in the pituitary during embryonic stages. Using zebrafish embryos, we establish the temporal dynamics of the development of corticotrophs. We demonstrate that embryonic perturbations in cortisol levels using chemical and environmental challenges affects the number of corticotrophs. The effect was greater at earlier stages of development but could be reversed. Finally, we demonstrate that early-life changes in glucocorticoid levels affect stress-related behaviours.

## Materials and Methods

### Zebrafish husbandry and maintenance

Zebrafish were maintained and bred according to FELASA guidelines (11) and all procedures were approved by the Institutional Animal Care and Use Committee of the Weizmann Institute, Israel. Adult fish were maintained in mixed sex groups at 28 C, pH 7 and conductivity 1000 µS/cm. Embryos were incubated at 28 C in 0.3X Danieau’s medium (17.4 mM NaCl, 0.21 mM KCl, 0.12 mM MgSO4, 0.18 mM Ca(NO_3_)_2_, 1.5 mM HEPES, pH 7.4) till 6 dpf. Transgenic zebrafish lines *pomc*:EGFP (12), *pomc*:Gal4 and Tg(*UAS*:BotxLCB-EGFP) (13) were used in this study.

### Chemical and genetic perturbations

Larvae were transferred to the wells of a 6-well plate in 3 ml of Danieau’s medium and treated with 0.2 mM aminoglutethimide or 35 µM dexamethasone (or left untreated as control). The medium and treatment were renewed every day; larvae were maintained at 28 C.

Embryos from a cross of *pomc*:Gal4 and *UAS:*BotxLCB-EGFP fish were sorted at 3 dpf based on the presence/absence of GFP in POMC cells in the pituitary.

### Imaging and image analysis

6 dpf larvae were euthanized by rapid chilling (2° to 4°C) and thereafter incubated in 4% paraformaldehyde overnight at 4 C. Larvae were washed with PBST (3 × 5 minutes each), transferred sequentially through 25%, 50% and 75% glycerol. Posterior part of the body, yolk sac and jaws were removed using insulin syringes. Larvae were placed ventral side facing the coverslip in 75% glycerol and mounted.

Imaging of samples was performed using Zeiss LSM800 confocal microscope using the oil immersion 40X objective. Imaging conditions, including laser intensities were kept constant between samples and groups throughout each experiment. Image J was used for analysis and cell counting. Specifically, the Cell Counter plug-in was used for counting after manual annotation of the first and last slices. Images depicted were generated by whole Z-stacks maximum intensity projections via ImageJ. Volocity was used for *pomc:gfp* intensity measurements (Find objects tool). Threshold for detection of signal and object size filter were kept constant for all samples (thresholding was based on SD of fluorescence ≥ 2; size of 0.5 to 50 μm^3^). Mean intensities of the clusters were normalized to their volume.

### Stress-recovery assay

The assay was performed as described earlier (14). Briefly, 6 dpf larvae were transferred to a well with 50% artificial seawater or fresh Danieau’s medium for 20 minutes. Following this, they were individually transferred to single wells of a 96-well plate with fresh Danieau’s medium. Locomotive behaviour of the larvae was monitored for 10 minutes in the dark using the Daniovision observation chamber (Noldus) to analyze recovery dynamics. Data acquired was analyzed using Ethovision XT12 (Noldus).

### Light-dark preference test

Experiments were performed between 10 a.m and 3 p.m. in a room with regular ambient light. Larvae were allowed to acclimate to the room prior to the assay. The light/dark apparatus consisted of a rectangular arena (5 cm X 3.5 cm X 1 cm), where one half is made dark by placing an infrared long-pass filter (Edmund Optics) over it. The arena was back-lit by infrared lighting facilitating uniform tracking in the entire arena and filled with 5 ml fresh Danieau’s medium and one larva was introduced into the lit half of the arena. Recording was begun prior to introducing the larva and the behaviour was monitored for 10 minutes using a near infrared sensitive camera (IO industries). Videos were analyzed using Ethovision XT12 (Noldus).

### Statistical analysis

Data are presented as mean ± standard error of mean (SEM). Analyses were performed using PRISM 6.0 (GraphPad Software Inc, San Diego, USA). Normality of the data was tested by the Shapiro-Wilk test. Depending on the outcome, parametric (ANOVA) or non-parametric tests (Mann-Whitney/Kruskal-Wallis test) were used for analysis. Post-hoc Sidak’s multiple comparisons test was performed when required.

## Results

### Changes in glucocorticoid levels during development impacts the corticotroph clusters

The optical transparency of zebrafish embryos makes them amenable to *in vivo* imaging of cellular components of the HPA axis. We used a reporter line, Tg(*pomc:*EGFP) (12), to monitor and quantify the number of corticotrophs during zebrafish development. Expression of *pomc:*EGFP was detected as early as 2 days post fertilization (dpf) during development (**Figure 1A**). Expression levels increased with time and by 4 dpf, two distinct anterior and posterior *pomc:*EGFP-positive clusters of cells were clearly visible in the developing pituitary (**Figure 1A**). Quantification of the number and the corresponding fluorescence intensity of GFP-positive cells in each cell cluster showed a steady time-dependent increase in both, the anterior and posterior pituitary (**Figures 1B, C, E, F**; *p<0*.*0001*, one-way ANOVA for analysis of **B, E**; for linear regression **C, F**: anterior cell number: *F=9*.*93, p=0*.*003*, posterior cell number: *F=87*.*71, p<0*.*0001*; anterior intensity: *F=27*.*86, p<0*.*0001*, posterior intensity: *F=30*.*64, p<0*.*0001*; n=11 larvae per group). The number and GFP intensity of cells in the anterior and posterior clusters was approximately similar and the ratio did not vary with time (**Figures 1D, G**; *p=0*.*0712* for cell number ratio, p=0.07082 for intensity ratio, one-way ANOVA), indicating that the size of the clusters maintained a steady state equilibrium. Though intensity and cell number measurements resulted in similar conclusions, we used intensity measurements for subsequent analysis since intensity measurements were more robust than cell number measurements at low levels of POMC expression.

**Figure 1:**
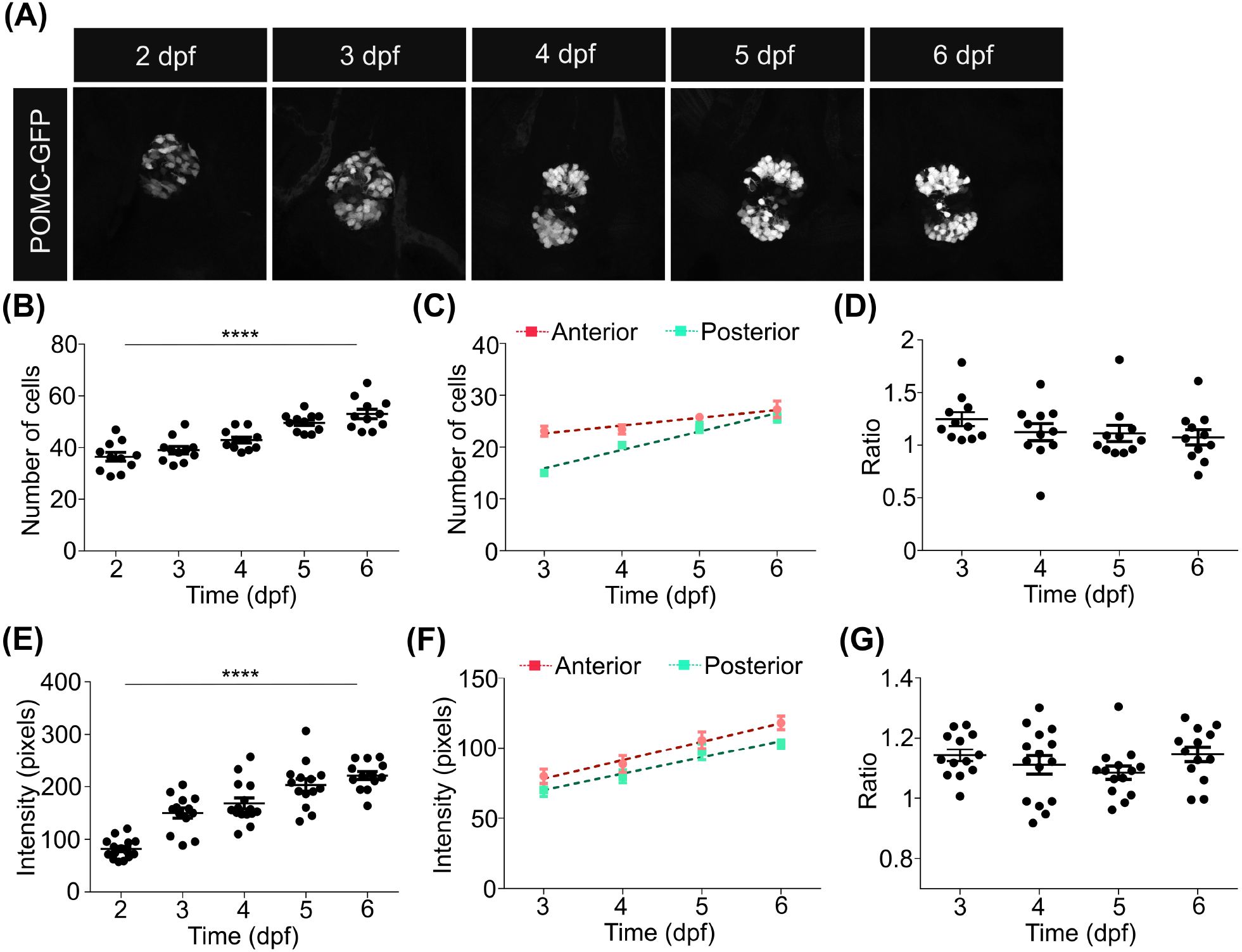
POMC cell number increases gradually during development. (**A**) Representative images from zebrafish larvae expressing GFP under a POMC promoter from 2 to 6 dpf. Larvae were collected and imaged every day to analyze the development of POMC positive cell clusters in the pituitary. n=11-15 per group (**B, E**) From the imaging performed in (A), the total number of GFP-positive cells (B) and the intensity of the GFP-positive clusters (E) were counted. (**C, D**) The number of cells in the anterior and posterior POMC-expressing clusters in the pituitary was quantified from day 3, when the clusters were distinct. While the number of cells in individual clusters showed an increase with time (C), and the relative number of cells in the anterior vs. the posterior cluster did not change (D). (**F, G**) The intensity of GFP-positive POMC cells in the anterior and posterior clusters of the pituitary was quantified from day 3. GFP intensity in individual clusters showed the same pattern as the cell number. Data was analyzed using linear regression. Data presented as mean±SEM, **** p<0.0001.

It has been shown that the HPA axis is modulated via a negative feedback mechanism through which cortisol inhibits the secretion of CRH from the hypothalamus and ACTH in the pituitary (2, 3). Following the analysis of the development of the POMC cell clusters, we sought to examine if such feedback inhibition also regulates the development ACTH-producing cells during embryonic pituitary development. Towards this, we performed chemical perturbations to modulate glucocorticoid levels and analyzed the effect on POMC cells in the pituitary. Larvae that were treated with aminoglutethimide, an inhibitor of cortisol synthesis, displayed an increase in the intensity of *pomc:EGFP* cells (**Figures 2A, B**; *p<0*.*0001*, n=15 larvae per group, one-way ANOVA comparing anterior and posterior clusters independently by Sidak’s multiple comparisons test). In contrast, larvae which were treated with dexamethasone, a cortisol analogue, displayed a marked decrease in the intensity of POMC cells especially in the anterior POMC cluster which is known to predominantly produce ACTH (12) (**Figures 2C, D**; *p<0*.*0001*, n=16 larvae per group, one-way ANOVA comparing anterior and posterior clusters independently by Sidak’s multiple comparisons test).

**Figure 2:**
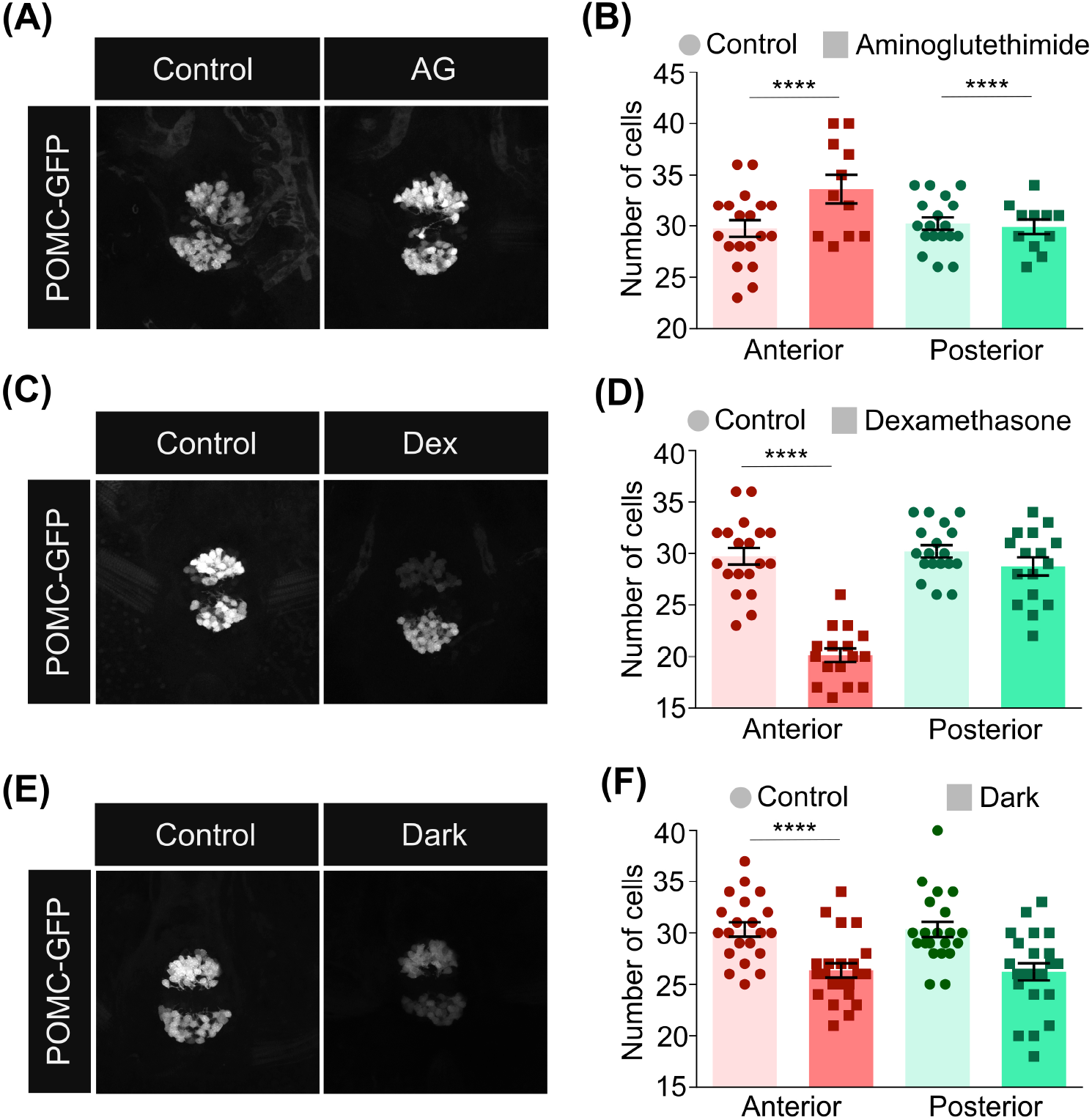
Perturbations of cortisol levels cause alterations in POMC cell clusters. (**A, B**) Larvae were treated with 0.2 mM aminoglutethimide, a cortisol synthesis inhibitor, from 3 to 5 dpf. Treated and untreated control larvae were collected at 6 dpf and fixed. The pituitary was imaged (A) and the intensity of POMC-positive cells in the anterior and posterior clusters was analyzed (B). n=19 control / 11 aminoglutethimide treated (**C, D**) Larvae were treated with 35 µM dexamethasone, a cortisol analogue, from 1 to 5 dpf. Treated and untreated control larvae were collected at 6 dpf and fixed. The pituitary was imaged (C) and the intensity of POMC-positive cells in the anterior and posterior clusters was analyzed (D). n=15 per group (**E, F**) Larvae were grown in the absence of lighting from 1 dpf to 5 dpf. Treated and control larvae (grown in the presence of light) were collected and fixed. The pituitary was imaged (E) and the intensity of POMC-positive cells in the anterior and posterior clusters was analyzed (F). n=21 per group Data presented as mean±SEM, *** p<0.001, **** p<0.0001.

We next examined whether corticotrophs are also affected by environmental stressors. To demonstrate this, we subjected the developing embryos to a chronic stressful experience, namely raising the embryos in the dark between 1-6 dpf. We observed a marked decrease in the intensity of POMC cells (**Figures 2E, F**; p<0.0001, n=30 control/35 treated, one-way ANOVA comparing anterior and posterior clusters independently by Sidak’s multiple comparisons test). These results suggested that changes in glucocorticoid levels as well as chronic environmental stressors during embryonic stages influence the development of pituitary glucocorticoid-producing POMC cells.

### Development of corticotrophs is sensitive to glucocorticoid levels at a critical developmental time window

We next explored whether the effect of glucocorticoids on corticotroph cell clusters is restricted to a specific developmental time window. To address this, we treated zebrafish larvae with dexamethasone at discrete time intervals and analyzed the effect on the POMC cell clusters. We observed that treatment with dexamethasone on or before 3 dpf, but not from 4 dpf, led to a consistent decrease in the size of POMC anterior cluster (**Figures 3A, B**; *p<0*.*0001*, n=10-15 larvae per group, one-way ANOVA comparing anterior and posterior clusters independently by Sidak’s multiple comparisons test). The posterior POMC cluster did not show a significant change in cell number (**Figures 3A, C**). To examine whether the developmental effect of glucocorticoids on corticotrophs is reversible, we treated larvae with dexamethasone between 1-6 days, and collected larvae at 6 hour intervals after withdrawing from the hormone followed by quantification of *pomc:*EGFP levels. This analysis showed that the decrease in POMC cells in the anterior cluster began to recover at 24 hours post-dexamethasone withdrawal and was complete by 48 hours from the time of withdrawal (**Figure 3D-F**; *p<0*.*0001*, n=10-16 larvae per group, one-way ANOVA followed by Sidak’s multiple comparisons test).

**Figure 3:**
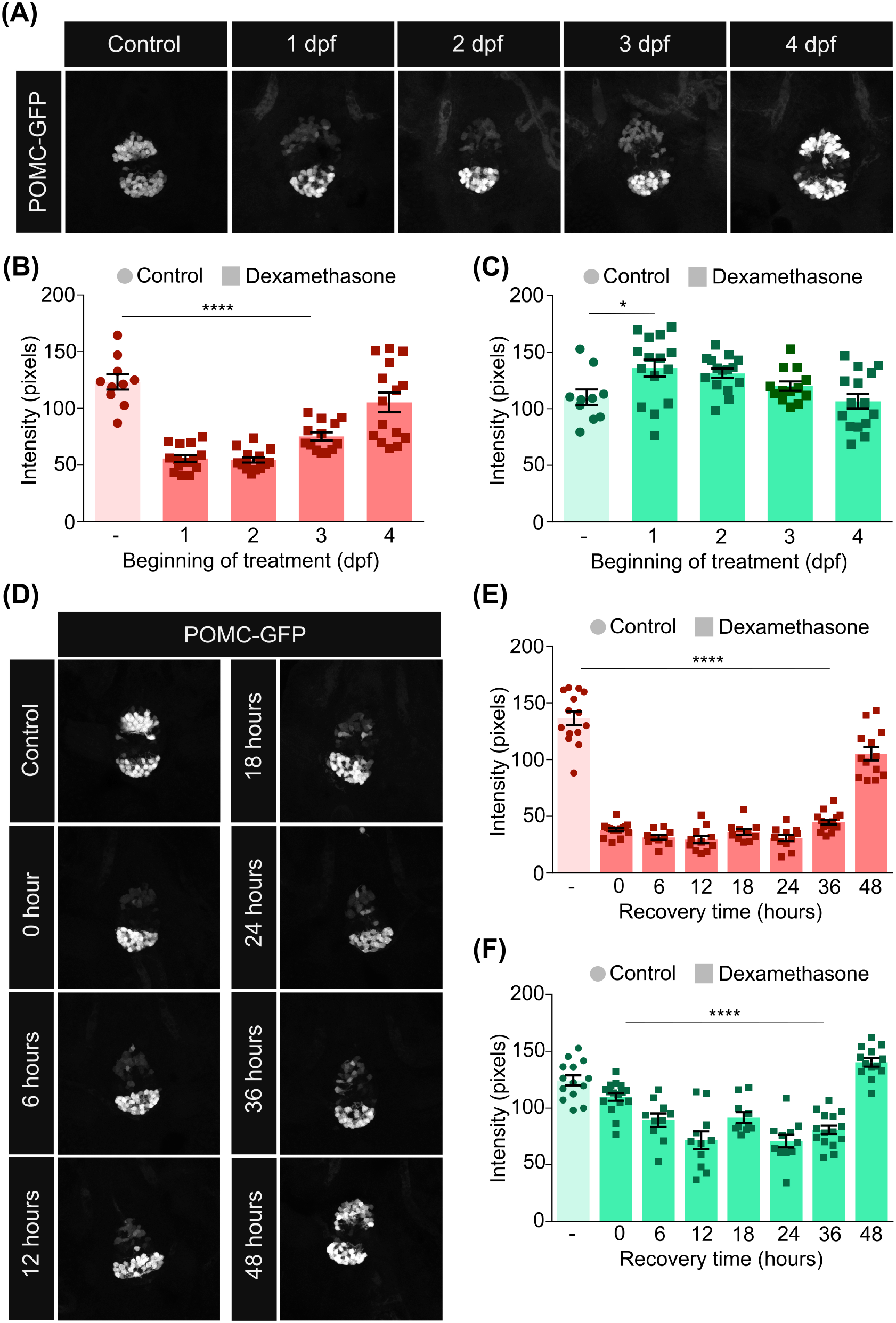
The effect of dexamethasone on POMC cells is drastic at early stages of development but reversible. (**A**) Larvae were treated with 35 µM dexamethasone beginning at 1, 2, 3 or 4 dpf. Treated and untreated control larvae were collected at 5 dpf, fixed and the pituitary was imaged. N=10-15 per group. (**B, C**) The intensity of GFP-positive POMC cells in the anterior and posterior clusters of the pituitary was quantified following different treatments of dexamethasone from (A). The effect of dexamethasone was more severe in the anterior cluster. (**D**) Larvae were treated with 25 µM dexamethasone from 1 dpf till 7 dpf. Dexamethasone was washed off from the larvae from 5 dpf at regular intervals. Larvae were collected at 7 dpf, fixed and the pituitary was imaged. N=11-14. (**E, F**) The intensity of GFP-positive POMC cells in the anterior and posterior clusters of the pituitary following different time of withdrawal from dexamethasone was quantified from (D). The reduction in the anterior cluster required 48 hours to come back to normal levels. Data presented as mean±SEM, *p<0.05, **** p<0.0001.

These results indicate that pituitary corticotrophs are particularly sensitive to increased glucocorticoid levels at early stages of development, but this effect can be reversed.

### Dysregulated glucocorticoid levels impact stress-related behaviour

We hypothesized that early-life perturbations of the feedback inhibition of POMC cells might impact stress-related behaviour. For this, we subjected dexamethasone treated larvae to an osmotic stressful challenge that induces freezing behaviour, a direct indicator of stress (14, 15). Upon treatment with dexamethasone, we observed a drastic impairment in the ability of larvae to recover from the stressful challenge (**Figure 4A**; *p<0*.*01* comparing control and dexamethasone following stress, n=8 unstressed, n=32 stressed, 2-way ANOVA followed by Sidak’s post-hoc multiple comparisons test). To further substantiate our claim, we used an additional zebrafish larval behavioural assay: the light-dark preference test, which is a standard measure of stress-responsive behaviour (9, 16, 17). In this assay, a freely swimming larva is introduced in an arena divided into a lit half and a dark half and its preference for each of the two halves is monitored. We observed that compared to control larvae, dexamethasone treated larvae spent more time in the dark zone (**Figure 4B**; *p=0*.*0486*, n=28 control/21 dexamethasone, Mann-Whitney test), indicating impaired stress-responsive behaviour.

**Figure 4:**
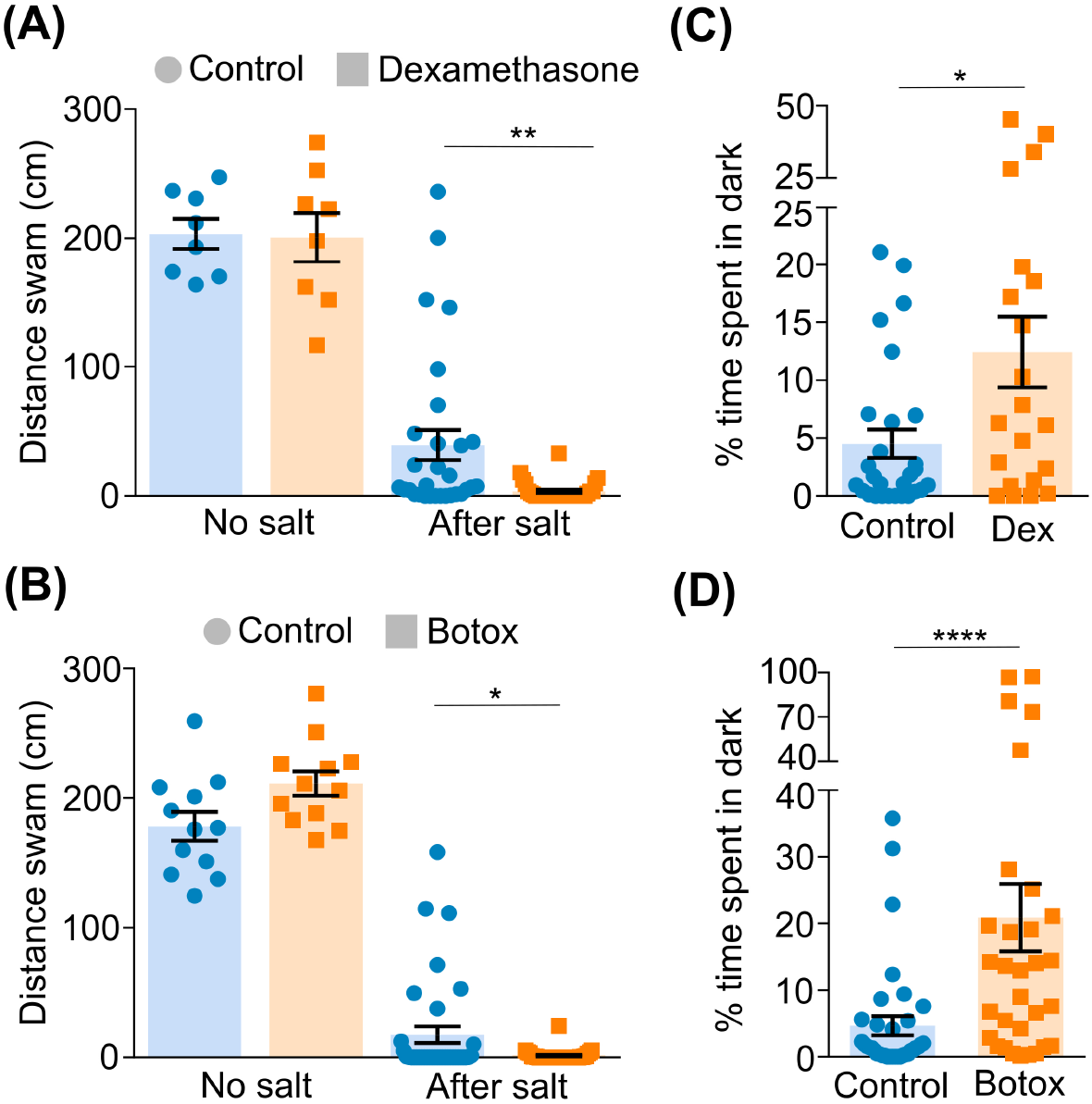
Altered ACTH levels result in impaired recovery from stress and anxiety-like behaviour. **(A, C**) Larvae were treated between 4-5 dpf with 35 µM dexamethasone/left untreated (A) or *Tg(pomc:Gal4;UAS:BoTxLCB-GFP)* were selected (C). At 6 dpf, control and treated/transgenic groups were subjected to osmotic stress by transferring to 50% artificial seawater for 20 minutes. Larvae were washed, transferred into individual wells of a 96-well plate, and their locomotive behaviour was recorded for 10 minutes to monitor recovery from the osmotic challenge. Behaviour of unstressed larvae was used as reference. (**B, D**) Larvae were treated with 35 µM dexamethasone between 4-5 dpf/maintained as control (B) or *Tg(pomc:Gal4;UAS:BoTxLCB-GFP)* larvae were selected (D). At 6 dpf, control and treated/transgenic larvae were introduced into the dark-light preference chamber and their behaviour was recorded for 10 minutes. The percentage of total time that the larvae spend in the dark zone was plotted. Data presented as mean±SEM, *p<0.05, **p<0.01, **** p<0.0001.

In order to confirm if the phenotype is specifically due to lower levels of circulating ACTH, we used a transgenic line *Tg(pomc:Gal4; UAS:BoTxLCB-GFP)*, where botulinum toxin light chain B is expressed specifically by corticotrophs, resulting in hampered synaptic release of ACTH-containing vesicles. When BoTxLCB-expressing larvae were subjected to osmotic stress, it led to a similar impaired recovery from stress (**Figure 4C**; *p=0*.*0007*, n=12 unstressed, n=36 stressed, 2-way ANOVA followed by Sidak’s post-hoc multiple comparisons test). An increase in exploration of the dark zone was also observed in the dark-light preference test in BoTxLCB larvae (**Figure 4D**; *p<0*.*0001*, n=36 control/32 Botox, Mann-Whitney test). These results show that altered corticotroph number and function result in impaired stress-responsive behaviour.

Taken together, our results show that pituitary corticotrophs cells appear early during development and are sensitive to changes in glucocorticoid levels at a critical developmental stages, with direct consequences on stress-responsive behaviour.

## Discussion

The HPA axis plays a crucial role in recovering from stressful situations, and begins to function early during development. However, how different players interact and influence the development of others is not known. Zebrafish embryos develop externally, and their optical transparency makes them amenable to the study of developmental phenomena. Using zebrafish embryos, we focused on recording the fine dynamics of the establishment of corticotrophs in the pituitary and the effect of fluctuations in cortisol levels on their development.

We observed that corticotrophs in the pituitary appear early during development, concurrent with previous studies showing that corticotrophs appear at 20 hpf (12, 18). Since the developmental dynamics of their establishment was not well-characterized earlier, we periodically monitored the development of the corticotrophs till 6 dpf. The corticotrophs appear as one mass between 24-48 hpf, which separates in two by 72 hpf, and their number increases steadily during development. However, the relative number of cells in the two clusters was unchanged with increase in cell number.

Cortisol levels are known to have a direct impact on the synthesis and release of ACTH from corticotrophs. However, whether this feedback regulation functions early during development and its effect on the development of corticotrophs is not well-characterized. Here, we demonstrate using a series of chemical and environmental perturbations that alterations in cortisol levels affect the development of corticotrophs from early stages. Pituitary corticotroph cells, especially the anterior ACTH-producing cluster were particularly sensitive to dexamethasone treatment during a critical early developmental window. Dexamethasone treatment earlier than 72 hpf had the most dramatic effect on the number of ACTH-positive cells in the anterior POMC cluster, coinciding with the developmental window when zebrafish larvae begin to show stress response by elevating cortisol levels at 3-4 dpf (7, 19, 20). This intense effect of dexamethasone treatment at early stages could be due to the HPA axis being in nascent stages of development.

We demonstrate the effect of fluctuations in cortisol levels on corticotrophs, primarily using cell cluster size as the output. Most previous studies have analyzed hormone synthesis and release, likely due to limited accessibility of corticotrophs in mammals. Hence, our measurement of cell number reveals that, in addition to its effect on dynamics of hormone availability, cortisol can directly impact the size of corticotroph population in the pituitary. Given that zebrafish embryos are amenable to live imaging, and the availability of various genetic tools, it would be interesting to further study the dynamics and mechanisms by which the regulation functions in the HPA axis.

Early life stress has been studied extensively in mammals with relation to its effect on behavior and susceptibility to mood disorders later in life (21-23). However, there is limited knowledge of the mechanisms that translate early life stress to behavioural changes in adulthood. Treatment with dexamethasone during developmental stages may mimic maternal/early life stress in mammalian embryos, and we show here that it can affect both, the development of corticotrophs and stress-responsive/anxiety-like behaviour. Our behavioural studies performed on zebrafish larvae show that developmental alterations in glucocorticoids levels as well environmental stressors are critical determinants that affect corticotrophs cell number and its associated stress coping behaviours.

## Author contributions

GP, AS and GL designed the study, GP and AS designed the experiments; GP performed the experiments and analyzed data with assistance from AS. AS and GL wrote the manuscript.

## Acknowledgements

We thank Roy Hofi, Estar Regev and the zebrafish facility personnel for animal care, Claire Wyart for the transgenic UAS:BoTxLCB-EGFP fish, and Uri Alon for discussions. G.L. lab is supported by the Israel Science Foundation (#349/21); US-Israel Bi-National Science Foundation (#2017325); Israel Ministry of Science (#3-16548); Hedda, Alberto, and David Milman Baron Center for Research on the Development of Neural Networks; Sagol Institute for Longevity Research; Maurice and Vivienne Wohl Biology Endowment and Foundation for Higher Education and Culture (FFHEC). G.L. is an incumbent of the Elias Sourasky Professorial Chair.

